# Susceptibility of *Pseudomonas aeruginosa* and *Acinetobacter baumannii* biofilms grown on denture acrylic to denture-cleaning agents and sonication

**DOI:** 10.1101/2022.09.21.508680

**Authors:** Marwah Behbehani, Ailbhe McDonald, Sean P Nair, Ingrid M Green

## Abstract

**Background:** Dentures can act as reservoirs for microorganisms that cause local and systemic diseases, including opportunistic pathogens *Pseudomonas aeruginosa* and *Acinetobacter spp*, leading causes of hospital-acquired pneumonia.

**Objectives:** To determine the efficacy of a brief cleaning protocol with four commonly used denture cleaning agents combined with ultra-sonic cleaning on *P. aeruginosa* and *A. baumannii* biofilms grown on denture acrylic. The effectiveness of a commercial sonicator on bacterial biofilm reduction was also evaluated.

**Materials and Methods:** *P. aeruginosa* and *A. baumannii* 48 h biofilms were grown in artificial saliva on polymethyl methacrylate (PMMA) discs, exposed to four disinfectants, with or without ultra-sonication, for 1 min and the number of viable bacteria post-exposure to the treatment determined. *P. aeruginosa* biofilms were exposed for one or five mins to two disinfectants plus a commercial sonicator.

**Results:** Ultra-sonication alone reduced total viable biofilm from 67.3% (*A. baumannii)* to 94.8% (*P. aeruginosa)*. Without ultra-sonication, 0.012% sodium hypochlorite, Steradent® Active Plus and Ecolab® 0.2% chlorhexidine reduced *P. aeruginosa* and *A. baumannii* biofilms viable counts by between 99.3% and >99.99%. Sonication further reduced 99.3% kill to undetectable levels (>99.99% kill). Colgate®Total Pro-Gum Health Daily mouthwash reduced *P. aeruginosa* by 17% and *A. baumannii* biofilms by 2.1 log. These number were further reduced by >99.99% with sonication (*p*=<0.001). Domestic sonicators, such as those for cleaning jewellery at home, were effective when used in conjunction with denture cleaning agents.

**Conclusion:** Rapid reduction of *P. aeruginosa* and *A. baumannii* biofilms on denture acrylic is possible with denture-cleaning agents and sonication

**Highlights:**

- Four antimicrobial agents tested against *P. aeruginosa* and *A. baumannii* biofilms
- 0.012% NaOCl and Steradent Active Plus were the most effective agents tested
- Sonication improved chlorhexidine and cetylpyridinium chloride-based agents

## Introduction

*Pseudomonas aeruginosa* and *Acinetobacter baumannii* are two important Gram-negative nosocomial pathogens, which pose an increasing challenge to healthcare providers ^1,2^. They are leading causes of hospital acquired pneumonia and bacteraemia ^1,3^, and are associated with increased mortality rates ^4^. *P. aeruginosa* and *A. baumannii* are adept at biofilm formation on skin, soft tissue and wounds, as well as abiotic surfaces including endotracheal tubes ^5,6^ Bacteria within biofilms are protected from hostile environments, such as antimicrobials, via a number of means such the biofilm neutralising antimicrobials by binding to them, the role of the exopolysaccharide matrices of a biofilm in limiting the rate of diffusion of the antimicrobial into the biofilm, and, as a consequence of oxygen deprivation within biofilms, a lower metabolic activity of bacteria can reduce the uptake of antimicrobial agents ^7^.

Dentures, worn by 20% of the total UK population ^8^, have been shown to harbour biofilms that can act as reservoirs of respiratory pathogens such as *P. aeruginosa* and *Acinetobacter spp* ^9,10^. *P. aeruginosa* and *A. baumannii* have secretion systems and efflux pumps capable of the removal of a wide variety of antibiotics, biocides detergents and antiseptics from the bacterial cell ^11,12^ which contributes to the challenge of biofilm removal. Moreover, *P. aeruginosa* has the ability to actively detach individual cells from its biofilm, a method called dispersal ^13^ implying that denture biofilms are a potential rich source for cross-infection, auto-infection and re-infection ^14^, with potential serious implications, particularly when patients are hospitalised and intubated.

Due to the topography of denture fitting surfaces, there is evidence that the use of both mechanical and chemical methods of disinfection is required for adequate cleaning and the removal of pathogenic or opportunistic bacteria from dentures ^15^. Mechanical cleaning with an ultrasonic device can be useful for physically and cognitively impaired patients ^16^ and sonication has been demonstrated to be an effective remover of Pseudomonas biofilms ^17^. However, laboratory grade sonicators are expensive, potentially hindering their use. Domestic sonicators, such as those used for home cleaning of jewellery, are more affordable and could be beneficial for denture cleaning both within hospitals and nursing homes, as well as for personal use.

The aim of this study was to determine whether commonly used denture cleaning agents, when combined with the mechanical effect of sonication, are effective at removing *P. aeruginosa* and *A. baumannii* biofilms grown on a commonly used denture material, polymethylmethacrylate (PMMA).

## Materials and Methods

### Denture Cleaning Agents and Sonication

*P. aeruginosa* (FR01) and *A. baumannii* (NCTC 1256) were grown in artificial saliva ^18^ at 37°C for 16 h with shaking at 225 rpm. The bacterial inoculums were diluted with artificial saliva to obtain an optical density (OD) of 0.05 at 600_nm_. Polymethylmethacrylate (PMMA) heat-cured denture resin discs (Lucitone 199®, Denstply, UK), 10 mm in diameter were fabricated (cured for 70°C for 6 h, 100°C for 4 h in a water-bath), and then polished (350-Grit, SiC Paper, Struers, UK; 500-Grit, Chaperline & Jacobs Ltd, Sutton, UK) to simulate the fit surface of the denture. Surface roughness (R_a_ μm) was verified by measuring 5 randomly selected points on 10 randomly selected discs, using a Proscan 1000 scanning laser profilometer (Scantron Industrial Products Ltd, England), using the displacement probe (KL131A) at 780_nm_ with a beam diameter of 15 - 35 μm and resolution of 0.02 μm. The R_a_ value was measured in both X and Y directions.

The discs were sterilised by autoclaving at 121.5°C for 15 min and positioned aseptically on light body silicone (Aquasil Ultra LV, Dentsply, USA) in sterile 12-well plates, exposing only the polished surface of the discs. To each well, 3 mL of the *P. aeruginosa* or the *A. baumannii* inoculum (OD 0.05) was added and incubated for 48 h at 37°C. Discs were aseptically removed after 48 h and dipped briefly in sterile PBS (Sigma-Aldrich, Merck Lifesciences Ltd, Dorset, UK), to remove any planktonic bacteria after which they were placed into 3 mL of one of the test agents: Steradent® Active Plus tablets (Reckitt Benckiser Ltd, Swindon, UK); 0.012% sodium hypochlorite (NaOCl) (Evans, Vanodine International PLC, UK); Ecolab® Chlorhexidine mouthwash (0.2% w/v Chlorhexidine digluconate) (CHX) (Ecolab® Ltd, Swindon, UK); Colgate® Total Pro Gum Health Daily Mouthwash, (0.075% w/v Cetylpyridinium chloride) (CPC) (Colgate-Palmolive, UK) or sterile phosphate-buffered saline (PBS). Using sterile distilled water, Steradent® Active Plus was prepared according to the manufacturer’s instructions and Evans bleach was diluted from the 4% w/v stock solution to achieve 0.012% w/v. Ecolab® CHX mouthwash and the Colgate® Total CPC mouthwash were used as supplied by the manufacturer. Half of the discs were sonicated in the active agents for 1 min using an ultra-sonic water-bath (Fisher Scientific, Germany), the remainder were not subjected to sonication but were exposed to the cleaning agents alone for 1 min. All discs were subsequently placed into 1 mL of neutralising buffer (Becton Dickinson, Detroit, MI, USA) and vortexed with glass beads (2.5 - 3.5 mm; VWR, Leicestershire, UK) for 1 min to disrupt the biofilm. The samples were serial diluted, the *A. baumannii* plated onto nutrient agar (Oxoid Ltd, Basingstoke, UK) and *P. aeruginosa* plated on to pseudomonas plates made with pseudomonas agar powder (Oxoid Ltd, Basingstoke, UK), supplemented with 1% glycerol (Sigma-Aldrich, Germany) and 1.2% agar technical (Oxoid Ltd, UK). The plates were incubated for 24 h at 37°C and the colonies counted allowing the number and percentage of bacteria killed by different antimicrobial agents and sonication regimes to be calculated.

### Domestic Sonicator Efficacy

The efficacy of a domestic sonicator (Ultra 7000, James Products, UK) in reducing viable biofilms, was compared to a laboratory ultra-sonic water-bath (FB 15047, Fisher Scientific, Germany) in conjunction with anti-microbial denture agents. To do this, *P. aeruginosa* 48 h biofilms were grown on PMMA discs as above. The discs were then removed aseptically and dipped briefly in sterile PBS to remove any planktonic bacteria prior to being immersed in 3 mL of one of the following agents: Ecolab® 0.2% CHX mouthwash, Colgate® Total Pro Mouthwash (CPC) or sterile PBS whilst being sonicated for 1 min or 5 min in either the laboratory ultrasonic water bath or the domestic sonicator. Control discs were subjected to the anti-microbial agents but not sonicated. The discs were then submersed in neutralising broth with glass beads, vortexed for 1 min and then serial diluted prior to being plated on to pseudomonas agar plates, and incubated for 24 h. The colonies were counted, and the number and percentage of bacteria killed was calculated. All experiments were performed in triplicate with three biological repeats.

### Statistical Analysis

Data were analysed using SPSS version 24 (SPSS Inc., Chicago, IL) using non-parametric tests, Kruskal-Wallis and Mann-Witney, to determine the number of viable bacteria post exposure to the antimicrobial test agents and sonication. The 5% level of statistical significance was used throughout the analyses.

## Results

### Surface roughness (Ra) and biofilm formation on PMMA discs

The arithmetic mean surface roughness of 10 acrylic discs was 5.2 μm (4.3 to 7.3 μm). See Fig 1A. *P. aeruginosa* and *A. baumannii* biofilms had an average CFU mL^-1^ per disc of 2.6 × 10^6^ and 3.87 × 10^5^ respectively. See Fig 1B.

**Figure 1.**
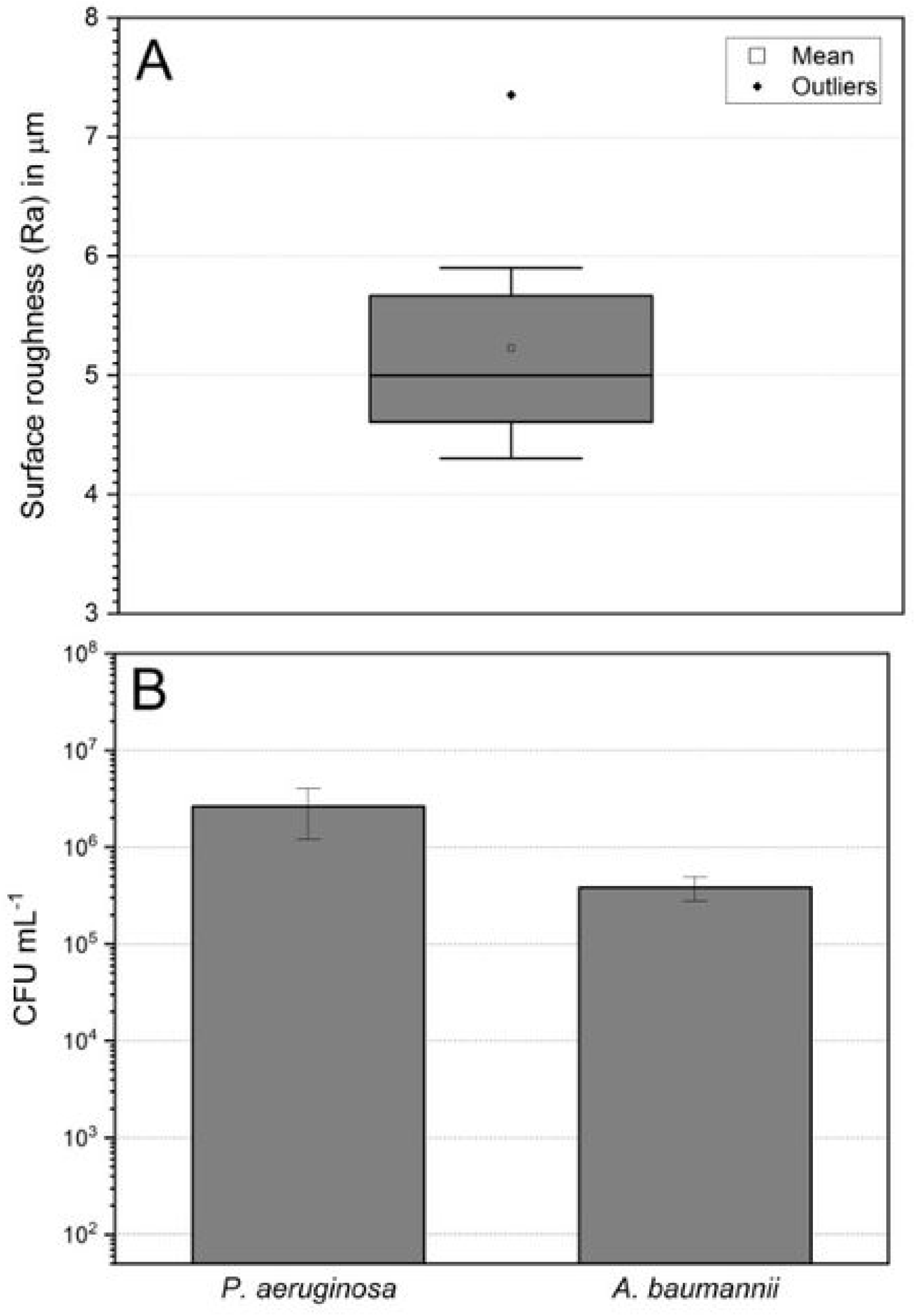
(A)Surface roughness (Ra) of 10 PMMA acrylic discs. The bold line represents the median, the box represents the 1^st^ and 3^rd^ quartiles, and the whiskers represent 95% of observations. (B) Colony forming units (CFU) per mL of 48 h P. aeruginosa and A. baumannii biofilms counted on 10 PMMA discs. Values represent the mean of 10 biofilms. Error bars are the ± SD.

### Susceptibility of biofilms to denture cleaning agents, with or without sonication

The 48 h biofilms were exposed to denture cleaning agents for 1 min. When compared to the PBS control, 0.012% NaOCl and Steradent® were the most effective anti-microbial agents evaluated, significantly reducing the median viable number of both *P. aeruginosa* and *A. baumannii* CFUs mL^-1^ by over 99.9% or 5.22 and 3.59 log respectively (p = <0.001). Ecolab® CHX mouthwash reduced viable *P. aeruginosa* biofilm numbers from 3.35 × 10^6^ to 1 × 10^4^ CFU mL^-1^ although this was not significant (p = 0.269), however it was able to significantly reduce *A. baumannii* viable counts from 7.95 × 10^4^ to <2 × 10^1^ CFU mL^-1^ (p = 0.001). Colgate® Total CPC mouthwash reduced the number of viable *P. aeruginosa* CFUs mL^-1^ by 17% (3.35 × 10^6^ to 1.5 × 10^6^ CFU mL^-1^) which was not statistically significant (*p* = 0.971), nor was its 2.1 log reduction of viable bacteria of the *A. baumannii* biofilms (*p*=0.945). See Fig 2.

**Figure 2.**
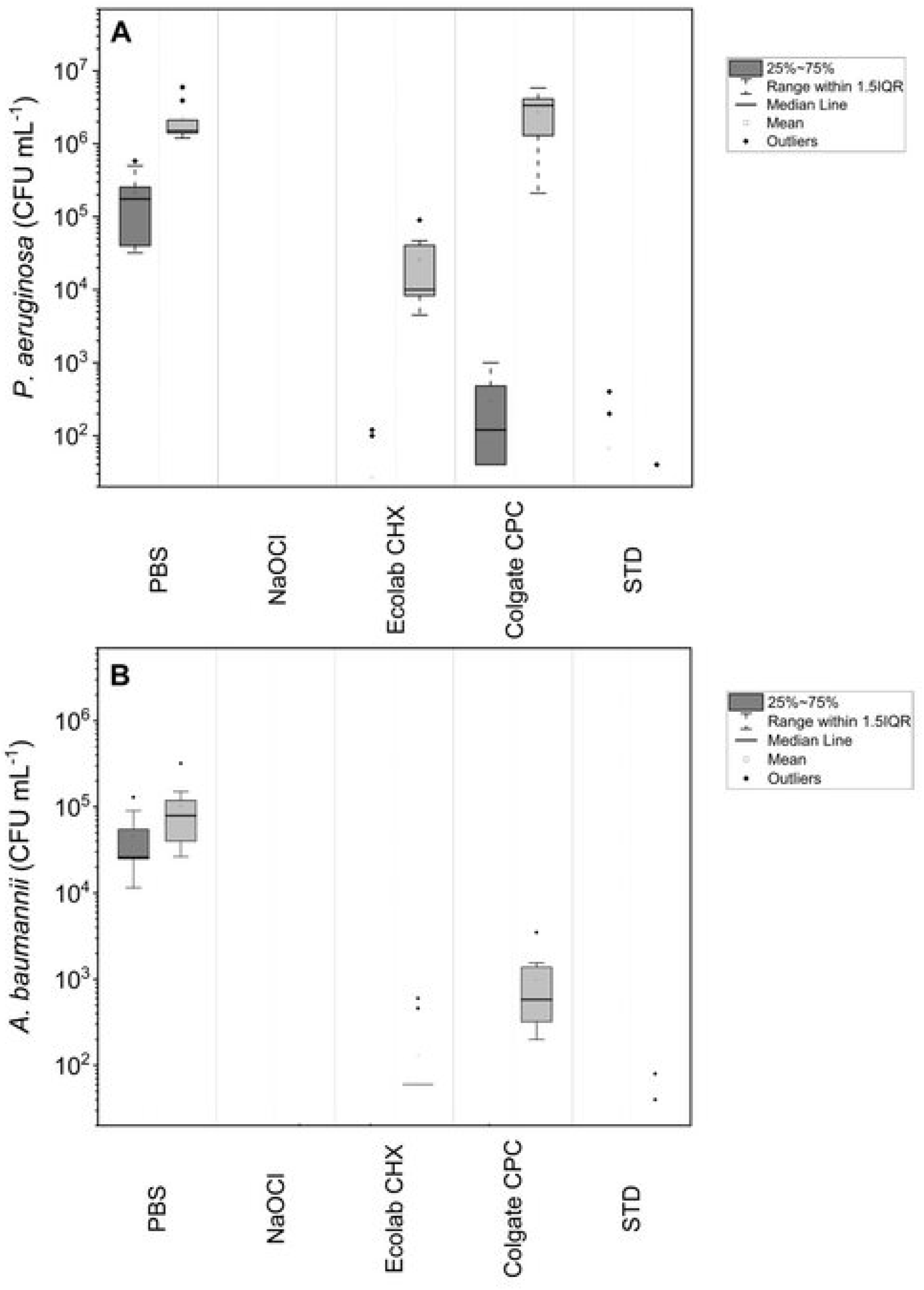
Survival of 48 h (A) *P. aeruginosa* and (B) *A. baumannii* biofilms after exposure to sodium hypochlorite (NaOCl), Ecolab chlorhexidine (Ecolab CHS), Colgate cetylpryidinium chloride (Colgate CPC), Steradent (STD) and PBS (control) with or without sonication, for 1 min. Light grey boxes represent exposure to denture cleaning agents only and dark grey boxes represent exposure to denture cleaning plus sonication. Each experiment was performed in triplicate with three biological repeats.

Sonication alone with the laboratory ultra-sonicator reduced biofilm viability counts by greater than 1 log for *P. aeruginosa* (p = <0.001) (3.35 × 10^6^ to 1.75 × 10^5^ CFU mL^-1^) and half a log for *A. baumannii* (7.95 × 10^4^ to 2.60 × 10^4^ CFU mL^-1^; p = 0.04). When sonication was used in conjunction with an antimicrobial, this significantly reduced the number of viable *P. aeruginosa* (3.33 × 10^6^ to 1.2 × 10^2^ CFU mL^-1^; p = <0.001) and *A. baumannii* CFU mL^-1^ (5.8 × 10^2^ to < 2 × 10^1^ CFU mL^-1^; p = <0.001) when compared to the cleaning agent only.

Sonication with Ecolab® CHX of *P. aeruginosa* biofilms led to a significant reduction in viable numbers compared to use of Ecolab® CHX only (1.0 × 10^4^ to <2 × 10^1^ CFU mL^-1^; p = <0.001). However no significant reduction was observed for *A. baumannii* biofilms as numbers were already very low with the Ecolab® CHX mouthwash alone (p = 0.258). Sonication did not significantly enhance NaOCl’s nor Steradent®’s kill rate for *P. aeruginosa* (p = 1.00; p = 0.666 respectively) or *A. baumannii* (p = 0.730; p = 0.436 respectively) biofilms as numbers were already very low or undetectable from use of the cleaning agent alone. See Fig 2.

On comparison of a domestic sonicator with the laboratory standard sonicator, there was no significant difference in the numbers of *P. aeruginosa* killed in biofilms when sonicated in PBS for either 1 min or 5 min (p = 0.190 and 0.113 respectively) by either machine. No difference between the sonicators could be determined with the Ecolab® CHX mouthwash as the number of viable CFU mL^-1^ were below the detection limit for both sonicators. When Colgate® Total CPC mouthwash was used in conjunction with 1 min sonication, the laboratory sonicator was significantly more effective than the domestic sonicator (p = <0.0001), however when the exposure time was increased to 5 min, no significant difference (p = 0.297) was observed. See Figure 3

**Figure 3.**
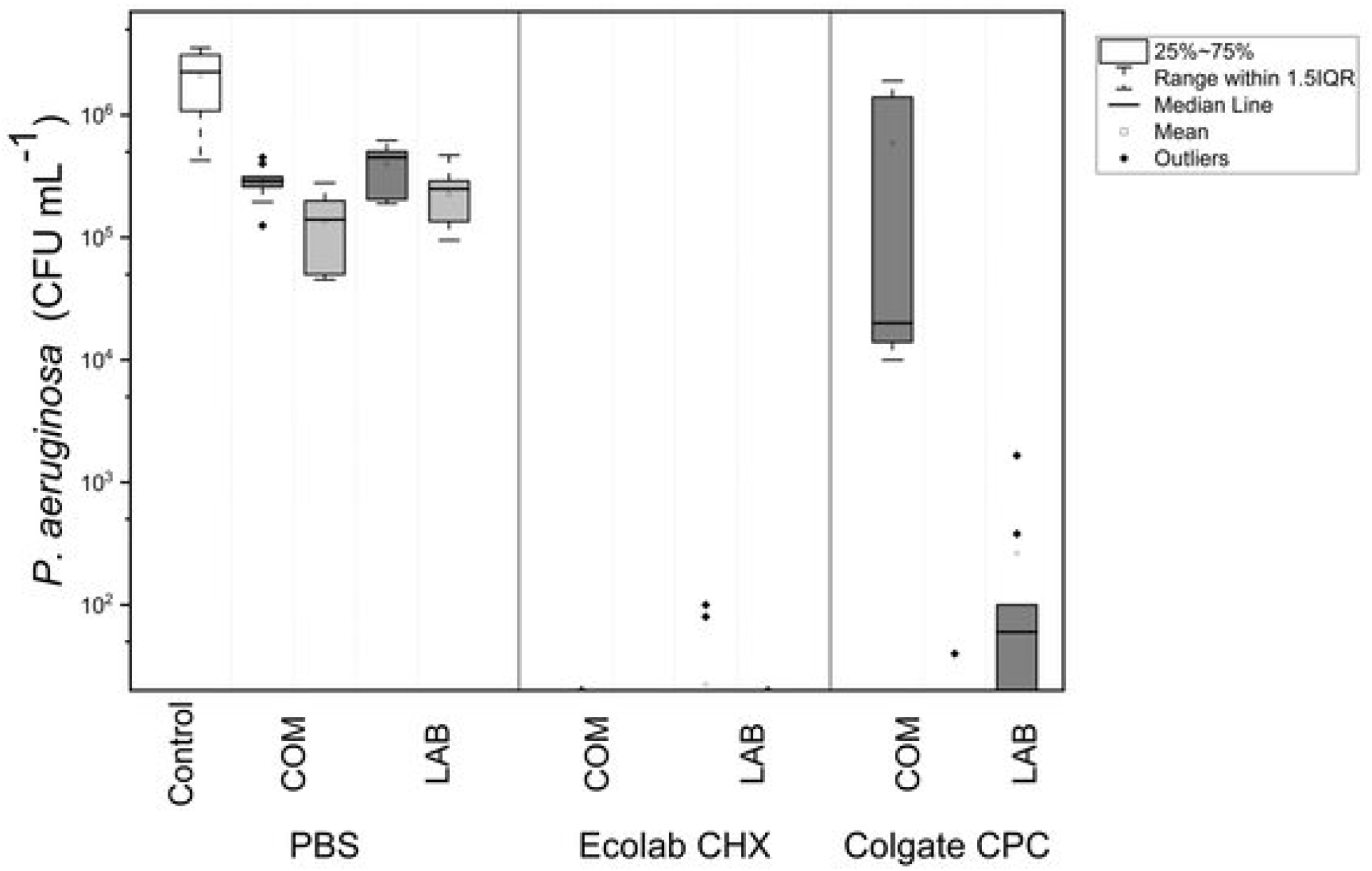
Survival of 48 h *P. aeruginosa* biofilms after exposure to Ecolab chlorhexidine (Ecolab CHS), Colgate cetylpryidinium chloride (Colgate CPC), and PBS (control) with sonication in either a commercial (COM) or laboratory (Lab) based sonicator for 1 min and 5 min. Dark grey boxes represent 1 min and light grey boxes represent 5 mins sonication times. The white box represents PBS without sonication as a control group. Each experiment was performed in triplicate with three biological repeats

## Discussion

*P. aeruginosa* and *A. baumannii*, potential respiratory pathogens that are proficient at biofilm formation, have been isolated from denture biofilms ^10^ and predispose denture users to aspirational pneumonia and systemic inflammation ^19^ Controlling denture plaque through an appropriate cleaning procedure is therefore important to reduce this risk ^20^.

A combination of chemical and mechanical denture cleaning methods is widely believed to be required for adequate denture cleaning ^15,21^, although there is a lack of consensus in the literature and from randomized controlled clinical trials on which of the available methods is best for disinfecting dentures ^15^. There is little data on the effectiveness of chemical disinfectants and/or ultrasonic cleaning against *P. aeruginosa* denture biofilms and no published data available on disinfection of *A. baumannii* biofilms from denture acrylic. This study aimed to determine how effective short exposure times to denture cleaning agents and sonication would be against *P. aeruginosa* and *A. baumannii* biofilms grown on denture acrylic, given that healthcare and social-care staff are under increased pressure to deliver care with limited resources. In addition, as the majority of denture wearers are elderly, often with reduced manual dexterity affecting their ability to brush their dentures thoroughly, a quick, accessible, and affordable ‘hands-off’ chemical and physical cleaning method would encourage perseverance with cleaning routines to attain thorough cleaning. This is particularly relevant, as up to 40% of denture wearers do not remove their dentures at night increasing the risk of pneumonia 2.3-fold when compared to those who do ^19^.

In this report, we tested four commonly used denture cleaning agents: 0.012% NaOCl, Steradent® Active Plus, Ecolab® 0.2% chlorhexidine mouthwash (CHX) and Colgate® Total Mouthwash 0.075% cetylpyridinium chloride (CPC) against 48 h *P. aeruginosa* and *A. baumannii* biofilms grown on denture acrylic resin with and without ultrasonic cleaning. NaOCl (0.012%) and Steradent® were very effective (p=<0.001) in reducing the numbers of both pathogens below the level of detection without sonication. Our results demonstrated that a short exposure time (1 min) to a 0.012% concentration of NaOCl led to significant reduction of viable CFU mL^-1^ of biofilms formed on denture acrylic resin, results that concurred with studies that used higher concentrations of NaOCl and longer exposure times. Salles et al. exposed *P. aeruginosa* biofilms grown for 48 h on acrylic resin to 0.25% and 0.5% w/v NaOCl for 20 min, resulting in undetectable CFU counts ^22^and Orsi et al. used even higher concentrations of NaOCl (1% and 2% w/v) for between 5 to 15 min on *P. aeruginosa* biofilms grown on acrylic resin, albeit with less mature (24 h) biofilms ^23^. Kohler et al. found that the *P. aeruginosa* MIC for NaOCl was significantly higher at 0.2% (w/v) than for *A. baumannii* (0.1% w/v) and demonstrated that *P. aeruginosa* was less susceptible to NaOCl than *A. baumannii* with 1 min contact time ^24^. However, as their work involved suspensions rather than biofilms, comparison with our study is difficult. Our results demonstrated that short exposure times of 1 min to a lower concentration of NaOCl (0.012%) led to a significant reduction of viable biofilms on denture acrylic resin.

Steradent® Active Plus showed comparable kills to 0.012% NaOCl against *P. aeruginosa* biofilms. This agrees with Mojarad et al. who showed that an alkaline peroxide (AP) based denture cleaner resulted in complete disinfection of acrylic dentures contaminated with *P. aeruginosa* 48 h biofilms, although their study had a significantly longer exposure time of 15 mins ^25^. Paranhos et al. also demonstrated that a 5 min exposure to an effervescent alkaline peroxide-based tablet was effective at significantly reducing *P. aeruginosa* CFU in 48 h biofilms grown on denture acrylic resin ^21^. Our study is the first to demonstrated that Steradent® Active Plus is capable of significantly reducing *A. baumannii* biofilm CFUs on denture acrylic PMMA.

The Ecolab® 0.25% chlorhexidine mouthwash reduced the viable count of the *A. baumannii* biofilms to levels comparable to 0.012% NaOCl and Steradent® but not when use against *P. aeruginosa* biofilms. This contrasts with a study by Masadeh et al, where they found that chlorhexidine mouthwashes, with and without ethanol, showed some activity against *P. aeruginosa* biofilms but no activity against *A. baumannii* biofilms that had been grown on polypropylene tubes ^26^. The high Mg2+ content in the outer membrane of *P. aeruginosa*, compared to other organisms, may confer intrinsic resistance of *P. aeruginosa* to chlorhexidine as chlorhexidine works by binding to negatively charged sites on the bacterial cell wall ^27^ and could explain the poor performance of these antimicrobial agents. The enhanced survival of *A. baumannii* has been demonstrated to be due to upregulation of genes for the major multi-drug efflux system, AdeAB, as well genes coding for an active chlorhexidine efflux protein, AceI, when *A. baumannii* is exposed to chlorhexidine ^28^.

Although, ethanol, an ingredient in Ecolab® 0.2% Chlorhexidine mouthwash, is known to be bactericidal against *P. aeruginosa* and *A. baumannii* in concentrations of greater than 30% v/v ^29,30^ it has been shown to increase the virulence of *A. baumannii* ^31^ and encourage biofilm formation in *P. aeruginosa* ^32^, particularly when present in lower concentrations. However, resistance of *P. aeruginosa* to antimicrobials has been related to several mechanisms, including enzymatic inhibition, efflux systems, alteration of outer membrane permeability, and genetic mutation ^27^.

The least effective agent in significantly reducing viable numbers of *P. aeruginosa* and *A. baumannii* biofilms when compared with the control group (PBS) was Colgate® Total Mouthwash, in which the active ingredient, CPC, an amphoteric surfactant, works in a similar way to chlorhexidine. Similar results were demonstrated by Masadeh et al who showed that a CPC based mouth wash exhibited poor activity against *A. baumannii* 48 h biofilms ^26^. There were no studies in the literature that have tested antimicrobial activity of CHX mouthwashes and CPC mouthwashes on *P. aeruginosa* or *A. baumannii* biofilms grown on dentures or denture materials such as PMMA.

Despite being effective antimicrobial agents, NaOCl and Alkaline Peroxide (AP), an active agent in Steradent, and CHX have been associated with several disadvantages, depending on the concentration used and treatment time of the denture materials. AP has also been shown to modify the colour of the acrylic significantly, whereas NaOCl has been shown to change the surface hardness and roughness, and noticeably alter the colour of the acrylic resin ^33,34^. A low NaOH concentration of 0.012% would have the benefit of reduced side effects whilst still efficiently disinfecting dentures, although it would be unsuitable for use on partial dentures, in which certain components are constructed from cobalt chrome. CHX, a frequently used antimicrobial agent and a common ingredient in mouthwashes, has been recognised as an increasing cause of anaphylaxis ^35^ and penetrates denture acrylic when used over a prolonged period of time ^36^. This causes staining of the denture materials but also, severe cytotoxicity to human gingival fibroblasts ^36^. A study by Röhner et al. compared 0.05% CHX with a 0.08% solution of NaOCl and found that not only was it less effective at eradicating *Staphylococcus aureus, S. epidermidis* and *Pseudomonas aeruginosa* biofilms from polyethlyene, but it also caused more cytotoxicity to human chondrocytes than NaOCl ^37^. In addition, an active ingredient of many mouthwashes, including Ecolab® 0.2% chlorhexidine, is ethanol, posing a hypothetical risk from repeated use of the development of oral cancer ^38^.

Ultrasonic cleaning is the use of high energy ultrasound waves in an aqueous solution to clean different objects. In this study, 1 min sonication with the control solution (PBS) was not as effective as chemical cleaning alone; however, when combined with Ecolab® CHX and Colgate® Total CPC mouthwashes, sonication significantly enhanced the ability of both agents to reduce viable counts (p = <0.0001). This has been supported by a previous study by Muqbil et al. which demonstrated the superiority of ultrasonic cleaning when combined with chemical cleaning compared to isolated treatment on *P. aeruginosa* biofilms ^39^. Our results support those of Nishi et al. who demonstrated the insufficiency of ultrasonic cleaning with water against Candida and bacterial spp. However, when combined with a chemical disinfectant, the resultant killing was higher than chemical cleaning alone ^40^, suggesting that it is not only the mechanical action of the ultrasonic waves that are responsible for killing but the combination of chemical and mechanical cleaning.

One of the limitations of laboratory or industrial sonicators is their cost. A less powerful domestic sonicator, such as those used for cleaning jewellery and personal appliances would be a more economical alternative, especially for personal home use. We tested a domestic sonicator, on its own and combined with Ecolab® CHX or Colgate® Total CPC mouthwashes, against *P. aeruginosa* biofilms. When combined with Ecolab® CHX there was no difference between the domestic and laboratory sonicator in the reduction of viable counts. However, with Colgate® Total CPC, a longer treatment time (5 min) was necessary to achieve a significant reduction in CFU ml^-1^. The domestic sonicator could therefore be an alternative to the laboratory sonicator when combined with chemical agents and longer treatment times, especially where loss of dexterity results in poor denture brushing and cleaning. Immersion in 0.12% NaOCl and Steradent® was statistically as effective as combined ultrasonic cleaning techniques, this indicates that the use of immersion alone with these denture cleaning agents is adequate for removal of *P. aeruginosa* and *A. baumannii* biofilms on denture acrylic.

## Conclusion

We have demonstrated that *P. aeruginosa* and *A. baumannii* biofilms are efficiently removed with low concentrations of NaOCl as well as Steradent® Active Plus and Ecolab® 0.2% CHX in conjunction with ultrasonic cleaning, in a short time frame of just 1 min. This regime could be particularly effective in an in-patient environment. With advancing age, a person’s sight and dexterity tend to deteriorate making brushing dentures challenging, thus the use of a domestic sonicator would aid denture cleaning in an increasingly elderly population. As certain individuals are reluctant to follow standard cleaning advice or remove their dentures overnight ^19^, this short efficient cleaning regime would be particularly useful.

## Conflict of interests

None declared.

## Ethical approval

Not required.

## Acknowledgments

This work was undertaken at University College London Hospitals/University College Hospital (UCLH/UCL), which received a proportion of funding from the Department of Health’s National Institute for Health Research (NIHR) Biomedical Research Centres Funding Scheme, UK.

## Abbreviations

CHX: chlorohexidine digluconate
NaOCl: sodium hypochlorite
AP: alkaline peroxide
CPC: cetylpyridinium chloride
PMMA: polymethylmethacrylate
PBS: phosphate-buffered saline
CFU: colony forming units

## Notes

### Competing Interest Statement

The authors have declared no competing interest.

### Summary of Updates

Abstract changed to structured, Figure 2 legends revised to clarify

